# Structural basis of SARS-CoV-2 Omicron immune evasion and receptor engagement

**DOI:** 10.1101/2021.12.28.474380

**Authors:** Matthew McCallum, Nadine Czudnochowski, Laura E. Rosen, Samantha K. Zepeda, John E. Bowen, Josh R. Dillen, Abigail E. Powell, Tristan I. Croll, Jay Nix, Herbert W. Virgin, Davide Corti, Gyorgy Snell, David Veesler

## Abstract

The SARS-CoV-2 Omicron variant of concern evades antibody mediated immunity with an unprecedented magnitude due to accumulation of numerous spike mutations. To understand the Omicron antigenic shift, we determined cryo-electron microscopy and X-ray crystal structures of the spike and RBD bound to the broadly neutralizing sarbecovirus monoclonal antibody (mAb) S309 (the parent mAb of sotrovimab) and to the human ACE2 receptor. We provide a structural framework for understanding the marked reduction of binding of all other therapeutic mAbs leading to dampened neutralizing activity. We reveal electrostatic remodeling of the interactions within the spike and those formed between the Omicron RBD and human ACE2, likely explaining enhanced affinity for the host receptor relative to the prototypic virus.

---

Although sequential COVID-19 waves have swept the world, no variants have accumulated mutations and mediated immune evasion to the extent observed for the SARS-CoV-2 Omicron (B.1.1.529) variant of concern (VOC). This VOC was first identified late November 2021 in South Africa and immediately designated a variant of concern (VOC) by the World Health Organization (*1*). Omicron has spread worldwide at an extraordinary pace compared to previous SARS-CoV-2 variants. The Omicron spike (S) glycoprotein, which promotes viral entry into cells (*2, 3*), harbors an unprecedented 37 residue mutations in the predominant haplotype as compared to the Wuhan-Hu-1 S or the 10 such substitutions in both the SARS-CoV-2 Alpha and Delta VOC (*4, 5*). The Omicron receptor-binding domain (RBD) and the N-terminal domain (NTD) contain 15 and 11 mutations, respectively, which lead to severe dampening of plasma neutralizing activity in infected or vaccinated individuals (*6*–*10*). Although the Omicron RBD harbors 15 residue mutations, it retains high affinity binding to human ACE2, the primary entry receptor, while gaining the capacity to recognize mouse ACE2 efficiently (*6, 11*). As a result of this antigenic shift, all authorized or approved therapeutic monoclonal antibodies (mAbs) lost their neutralizing activity against Omicron with the exception of S309 (sotrovimab parent) and the COV2-2196/COV2-2130 cocktail (cilgavimab/tixagevimab parent), which respectively experienced 2-3-fold and 12-200-fold reduced potency using pseudovirus or authentic virus assays (*6*–*10*). This extent of evasion of humoral responses has important consequences for therapy and prevention of both the current pandemic, and future pandemics. Defining the molecular mechanisms responsible for these facts is important.

To provide a structural framework for the observed Omicron immune evasion and receptor recognition, we determined cryoEM structures of the prefusion-stabilized SARS-CoV-2 Omicron S ectodomain trimer bound to S309 and S2L20 (NTD-specific mAb) Fab fragments **(Fig. 1 and fig. S1)** and the X-ray crystal structure of the Omicron RBD in complex with human ACE2 and the Fab fragments of S309 and S304 at 3.0 Å resolution. Surface plasmon resonance (SPR) based binding studies were used to determine binding affinities of mAbs to the Omicron RBD.

**Fig. 1.**
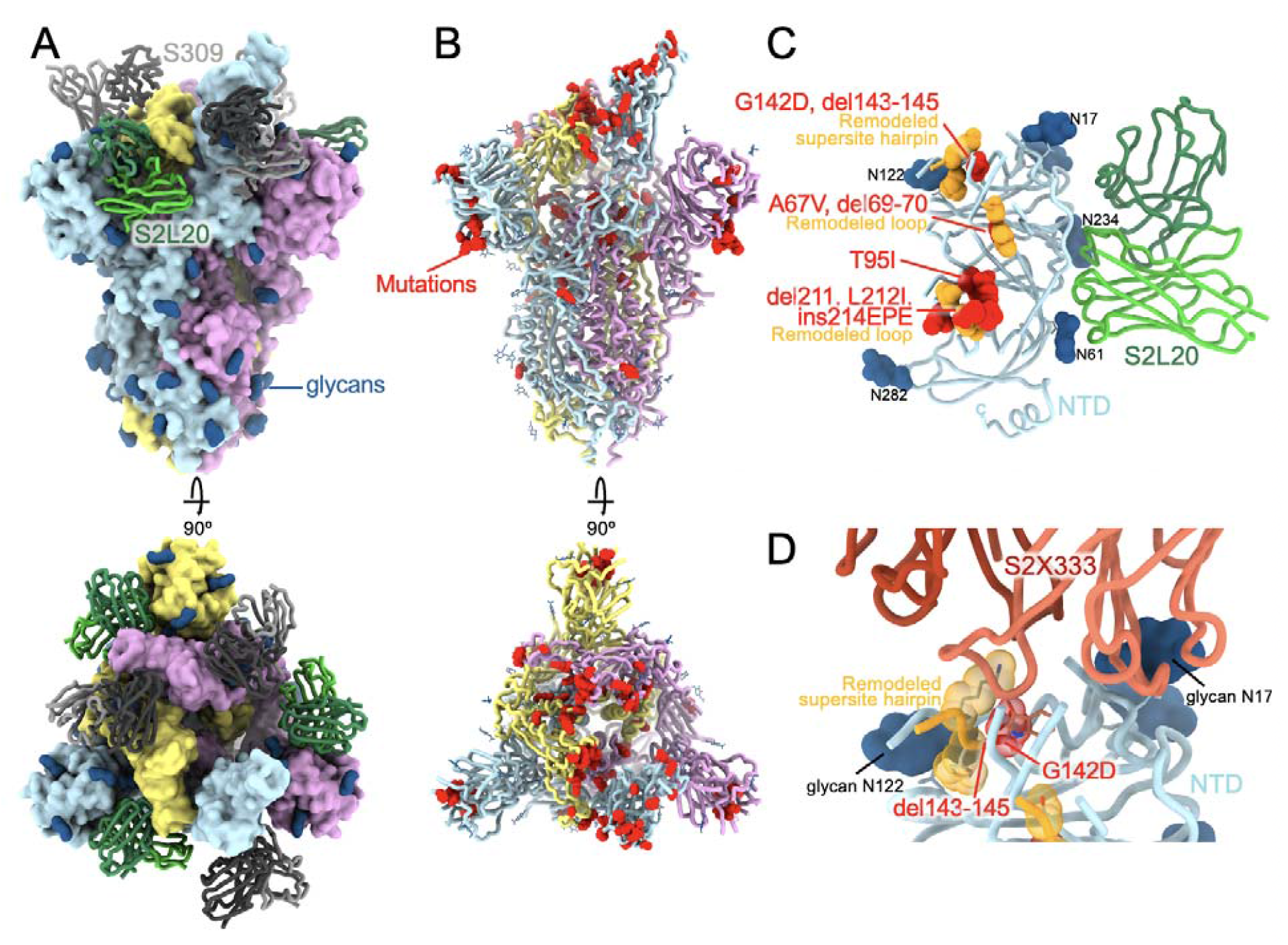
CryoEM structure of the SARS-CoV-2 Omicron S reveals a remodeling of the NTD antigenic supersite. **(A)** Surface rendering in two orthogonal orientations of the Omicron S trimer with one open RBD bound to the S309 (grey) and S2L20 (green) Fabs shown as ribbons. (**B**) Ribbon diagrams in two orthogonal orientations of the S trimer with one open RBD with residues mutated relative to Wuhan-Hu-1 shown as red spheres (except D614G which is not shown). In panels A-B, the three S protomers are colored light blue, pink or gold. (**C)** The S2L20-bound Omicron NTD with mutated, deleted or inserted residues rendered or indicated as red spheres. Segments with notable structural changes are shown in orange and labeled. (**D)** Zoomed-in view of the Omicron NTD antigenic supersite highlighting incompatibility with recognition by the S2×333 mAb (*15*) (used here as an example of prototypical NTD neutralizing mAb). N-linked glycans are shown as dark blue surfaces.

3D classification of the cryoEM data revealed the presence of two conformational states with one or two RBDs in the open conformation for which we determined structures at 3.2 Å and 3.1 Å resolution, respectively **(Fig. 1A-B and fig. S1)**. Merging the particles from both conformations and refining with C3 symmetry improved map resolution to 2.6 Å. Focused classification and local refinement of the S309-bound RBD and of the S2L20-bound NTD were used to account for their conformational dynamics and improve local resolution of these regions to 2.9 and 3.3 Å resolution, respectively.

Whereas most VOC have only a few mutations beyond the NTD, RBD, and furin cleavage site regions, the Omicron spike harbors eight substitutions outside of these areas: T547K, H655Y, N764K, D796Y, N856K, Q954H, N969K, and L981F, which could all be modeled in the map **(Fig. 1A-B and Fig 2)**. Four of these mutations introduce electrostatic contacts formed between core S_2_ subunit helices and the S_1_ subunit: N764K binds Q314, T547K binds S982, N856K binds D570, and N969K binds Q755 **(Fig. 2)**. Nearby, L981F is also localized to the core helices of S2 and improves hydrophobic packing **(Fig. 2)**. These mutations are next to the prefusion-stabilizing 2P mutations (K986P and V987P) used in all three vaccines deployed in the US **(Fig. 2)**. Enhanced interactions between the S_1_ and S_2_ subunits in Omicron S, along with altered processing at the S_1_/S_2_ cleavage site due to the N679K and P681H mutations, might reduce S_1_ shedding, consistent with recent studies (*12, 13*). Dampened S_1_ subunit shedding could enhance the effector function activity of vaccine- or infection-elicited Abs along with that of therapeutic mAbs that retain Fc-mediated effector function (*14*).

**Figure 2.**
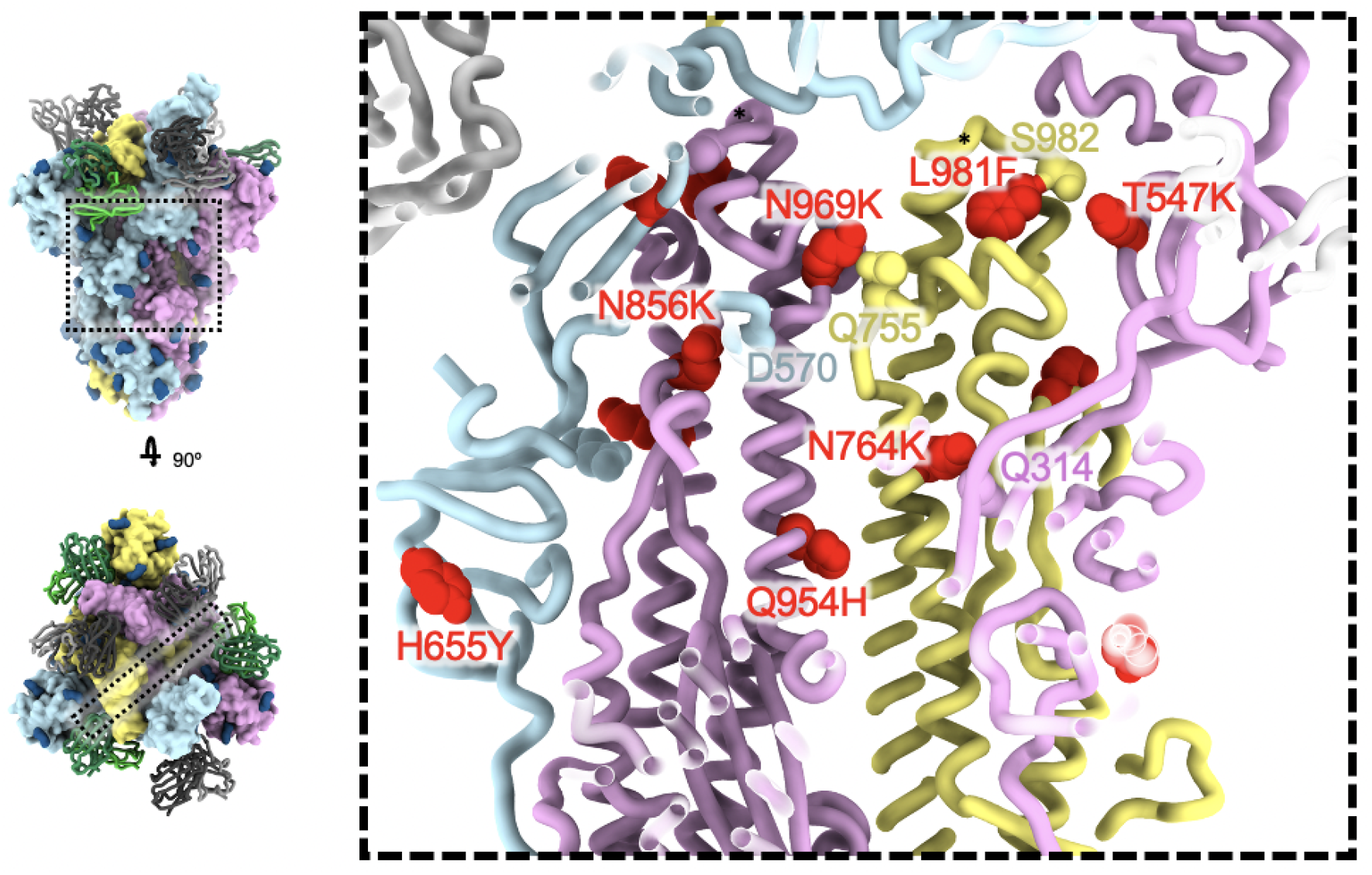
SARS-CoV-2 Omicron S fusion machinery mutations. A cross section through the core of the spike glycoprotein is shown (the location of this slice on the spike glycoprotein is shown on the left). Mutations T547K, H655Y, N764K, N856K, Q954H, N969K, and L981F are shown as red spheres; residues these mutations interact with are shown as spheres colored as the protomer they belong to. Black asterisks show the position of residues involved in the prefusion-stabilizing 2P mutations (K986P and V987P) used in all three vaccines deployed in the US.

The Omicron NTD carries numerous mutations, deletions (del), and an insertion (ins) including A67V, del69-70, T95I, G142D, del143-145, del211, L212I, and ins214EPE, which are all resolved in the cryoEM map **(Fig. 1C)**. Many of these mutations have been described in previously emerged VOC: del69-70 was found in Alpha, T95I was present in Kappa and Iota, whereas G142D was present in Kappa and Delta. 211del, L212I, and ins214EPE are adjacent to T95I, hinting that this site could have functional relevance, though these mutations are far from the NTD antigenic supersite and might not be involved in immune escape (*15*–*17*). G142D and del143-145 map to the NTD antigenic supersite (site i) beta-hairpin and alter its structural organization which would be incompatible with binding of the potent S2×333 NTD neutralizing mAb **(Fig. 1D)** (*4, 15*). Moreover, del143-145 is reminiscent of the Alpha del144 which was also isolated as an escape mutation in the hamster challenge model in presence of S2×333 leading to viral breakthrough (*15*). These data indicate that G142D and del143-145 account for the observed SARS-CoV-2 Omicron evasion from neutralization mediated by a panel of NTD mAbs (*6, 8*).

The RBD is the main target of plasma neutralizing activity in convalescent and vaccinated individuals and comprises several antigenic sites recognized by neutralizing Abs with a range of neutralization potencies and breadth (*14, 18*–*32*). Our structure resolves the complete RBD and provides a high-resolution blueprint of the residue substitutions found in this variant (**Fig. 3A**) and their impact in antibody binding (**Table 1**). The K417N, G446S, S477N, T478K, E484A, Q493R, G496S, Q498R, N501Y and Y505H mutations are part of antigenic site I, which was shown to be immunodominant (*18*). K417N, E484A and Q493R lead to loss of electrostatic interactions and steric clashes with REGN10933 whereas G446S would lead to steric clashes with REGN10987, as confirmed by the absence of detectable binding to the Omicron RBD (**Fig. 3 B-C and Table S1**). S477N, T478K and Q493R would sterically impair COV2-2196 recognition whereas E484A leads to loss of a hydrogen bond with COV2-2130 explaining the abrogated and markedly reduced binding of COV2-2196 and COV2-2130 to the Omicron RBD relative to the Wuhan-Hu-1 RBD, respectively (**Fig. 3 D-E and Table S1**). E484A leads to loss of hydrogen bonds with LY-CoV555 heavy and light chains and Q493R prevents binding through steric hindrance (**Fig. 2F and Table S1**). K417N is expected to negatively affect the constellation of electrostatic interactions formed between the Omicron RBD and LY-CoV16 heavy chain abolishing binding (**Fig 3H and Table S1**). Finally, K417N E484A and Q493R hinder CT-P59 engagement through a combination of steric hindrance and remodeling of electrostatic contacts, thereby preventing binding (**Fig. 3H and Table S1**). These data are consistent with previous analyzes of the effect of individual amino acid mutations using deep mutational scanning studies of yeast-displayed RBDs (*7, 27, 33*–*35*).

**Fig. 3.**
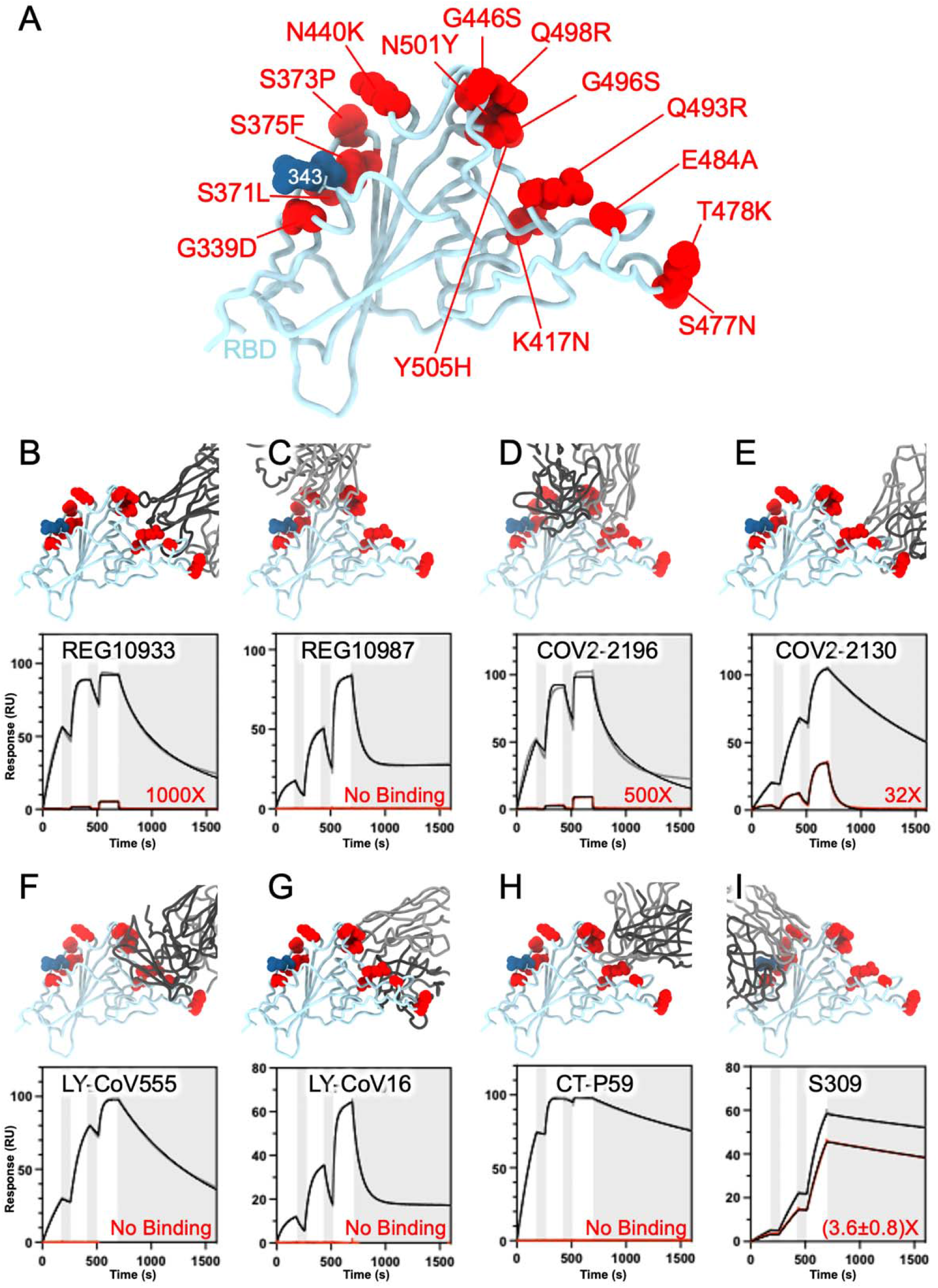
SARS-CoV-2 Omicron RBD mutations promote escape from a panel of clinical mAbs. **A**, Ribbon diagram of the RBD with residue mutated relative to the Wuhan-Hu-1 RBD shown as red spheres. The N343 glycan is rendered as blue spheres. **B-I**, Zoomed-in view of the Omicron RBD superimposed to structures of the RBD bound to REGN10933 (B), REGN10987 (C), COV2-2196 (D), COV2-2130 (E), LY-CoV555 (F), LY-CoV16 (G), CT-P59 (H) or S309 (I). Binding of the Wuhan-Hu-1 (gray line) or Omicron (red line) RBD to the corresponding mAb was evaluated using surface plasmon resonance (single-cycle kinetics) and is shown at the bottom. The black line is a fit to a kinetic model. The decrease in affinity between Wuhan-Hu-1 and Omicron binding is indicated in red.

**Table 1:**
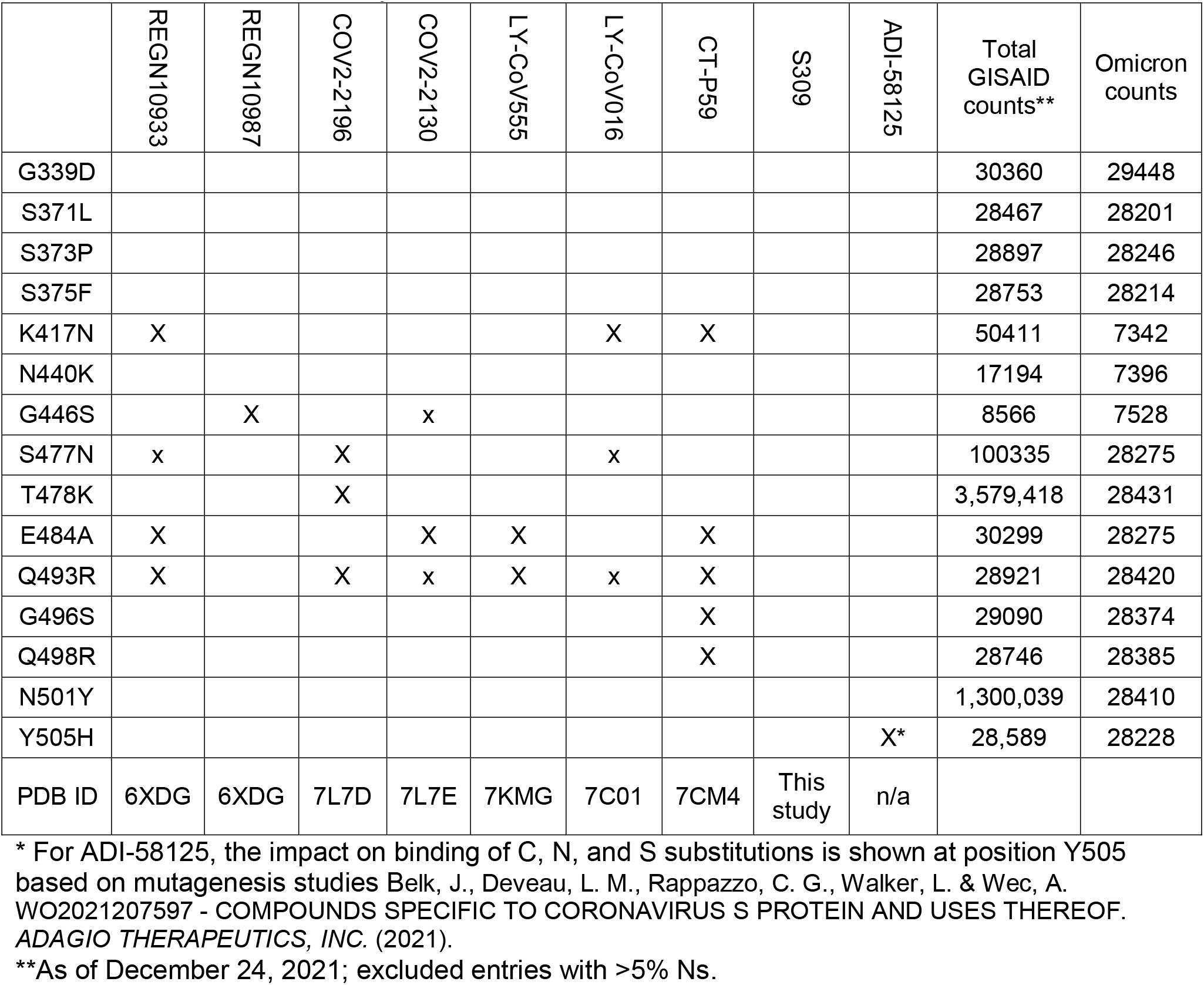
Omicron RBD mutations likely (X) or potentially (x) impacting the binding of therapeutic mAbs based on structural analyses.

|  | REGN10933 | REGN10987 | COV2-2196 | COV2-2130 | LY-CoV555 | LY-CoV016 | CT-P59 | S309 | ADI-58125 | Total GISAID counts** | Omicron counts |
| --- | --- | --- | --- | --- | --- | --- | --- | --- | --- | --- | --- |
| G339D |  |  |  |  |  |  |  |  |  | 30360 | 29448 |
| S371L |  |  |  |  |  |  |  |  |  | 28467 | 28201 |
| S373P |  |  |  |  |  |  |  |  |  | 28897 | 28246 |
| S375F |  |  |  |  |  |  |  |  |  | 28753 | 28214 |
| K417N | X |  |  |  |  | X | X |  |  | 50411 | 7342 |
| N440K |  |  |  |  |  |  |  |  |  | 17194 | 7396 |
| G446S |  | X |  | x |  |  |  |  |  | 8566 | 7528 |
| S477N | x |  | X |  |  | x |  |  |  | 100335 | 28275 |
| T478K |  |  | X |  |  |  |  |  |  | 3,579,418 | 28431 |
| E484A | X |  |  | X | X |  | X |  |  | 30299 | 28275 |
| Q493R | X |  | X | x | X | x | X |  |  | 28921 | 28420 |
| G496S |  |  |  |  |  |  | X |  |  | 29090 | 28374 |
| Q498R |  |  |  |  |  |  | X |  |  | 28746 | 28385 |
| N501Y |  |  |  |  |  |  |  |  |  | 1,300,039 | 28410 |
| Y505H |  |  |  |  |  |  |  |  | X* | 28,589 | 28228 |
| PDB ID | 6XDG | 6XDG | 7L7D | 7L7E | 7KMG | 7C01 | 7CM4 | This study | n/a |  |  |

\* For ADI-58125, the impact on binding of C, N, and S substitutions is shown at position Y505 based on mutagenesis studies Belk, J., Deveau, L. M., Rappazzo, C. G., Walker, L. & Wec, A. WO2021207597 - COMPOUNDS SPECIFIC TO CORONAVIRUS S PROTEIN AND USES THEREOF. ADAGIO THERAPEUTICS, INC. (2021).
\*\*As of December 24, 2021; excluded entries with >5% Ns.

The SARS-CoV-2 Omicron G339D and N440K mutations are within or nearby antigenic site IV, which is recognized by the S309 mAb (*20, 29*). S309 retains its neutralizing activity with a 2 to 3-fold reduced potency relative to that determined against Wuhan-Hu-1 pseudovirus or Washington-1 authentic virus (*6, 8*–*10*). The lysine side chain introduced by the N440K substitution points away from the S309 epitope and does not affect binding. The aspartic acid side chain introduced by the G339D substitution also points away from the S309 epitope, though it appears to reorient the N343 glycan (**fig S2**). The moderate reduction of the Omicron RBD binding to S309 (**Fig 3I and Table S1**) mirrors the 2-3-fold reduced neutralization potency of this VOC, relative to prototypical viruses, and concurs with DMS analysis of individual mutations on S309 recognition.

We recently reported that the SARS-CoV-2 Omicron RBD binds human ACE2 with a ∼2.4 fold enhanced affinity relative to the Wuhan-Hu-1 RBD (*6*). To understand how the constellation of RBD mutations present in the SARS-CoV-2 Omicron RBD impact receptor recognition, we determined a crystal structure of the SARS-CoV-2 Omicron RBD bound to the human ACE2 ectodomain along with the S309 and S304 Fab fragments at 3.0 Å resolution **(Fig. 4A and Table S2)**. The N501Y mutation alone is known to enhance ACE2 binding by a factor 6, as observed with the Alpha variant (*5*), likely as a result of increased shape complementarity between the introduced tyrosine side chain and the ACE2 Y41 and K353 side chains **(Fig. 4B)**. The K417N mutation is known to dampen receptor recognition ∼3-fold via loss of a salt bridge with ACE2 D30 (*4, 5, 36, 37*) **(Fig. 4C)**.The Q493R and Q498R remodel electrostatic interactions with ACE2 and introduce two new salt bridges **(Fig. 3D)**. Both of these individual mutations reduce ACE2 binding affinity slightly by DMS studies of the yeast-displayed SARS-CoV-2 RBD (*38*). Finally, S477N leads to formation of a new hydrogen bond between the introduced asparagine side chain and the ACE2 S19 side chain hydroxyl **(Fig. 3E)**. Collectively, these mutations have a net enhancing effect on binding of the Omicron RBD to human ACE2, relative to Wuhan-Hu-1, indicating of structural epistasis enabling immune evasion and retention of efficient receptor engagement. This is illustrated by the large number of Omicron mutations in the immunodominant receptor-binding motif, which likely explains a significant proportion of the loss of neutralization by convalescent and vaccine-elicited polyclonal antibodies, in line with the know plasticity of this subdomain.

**Figure 4.**
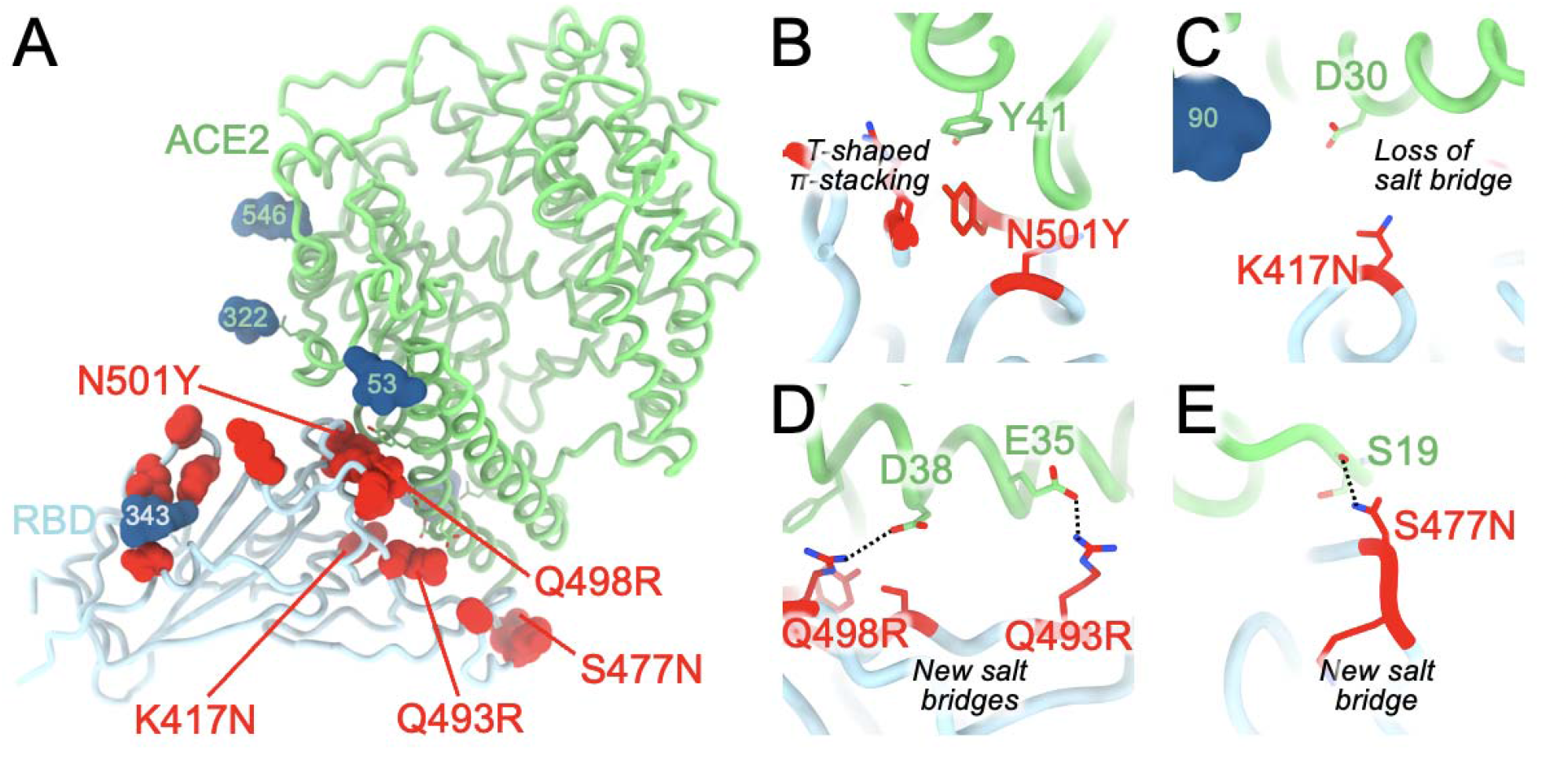
Molecular basis of human ACE2 recognition by the SARS-CoV-2 Omicron RBD. **A**, Ribbon diagram of the Omicron RBD in complex with the ACE2 ectodomain. The S309 and S304 Fab fragments are not shown for clarity. **B-E**, Zoomed-in views of the RBD/ACE2 interface highlighting modulation of interactions as a result of introducing the N501Y (B), K417N (C), Q493R and Q498R (D) and S477N (E) residue substitutions.

Although the N501Y mutation has previously been described to enable some SARS-CoV-2 VOC to infect and replicate in mice, the Alpha and Beta variant RBDs only weakly bound mouse ACE2 (*39, 40*). The SARS-CoV-2 Omicron RBD, however, interacts more strongly with mouse ACE2 than the aforementioned variant RBDs when evaluated side-by-side **(fig S3 A)** and can utilize mouse ACE2 as an entry receptor for S-mediated entry (*6, 11*). We propose that the Q493R mutation plays a key role in enabling efficient mouse ACE2 binding, through formation of electrostatic interactions with the N31 side chain amide (K31 in human ACE2), as supported by in silico modeling based on our human ACE2-bound crystal structure **(fig S3 B)**. These findings concur with the emergence and fixation of the Q493K RBD mutation upon serial passaging in mice to yield a mouse-adapted virus designated SARS-CoV-2 MA10 (*41*).

This work defines the molecular basis for the broad evasion of humoral immunity exhibited by SARS-CoV-2 Omicron and underscores the SARS-CoV-2 S mutational plasticity and the importance of targeting conserved epitopes for vaccine and therapeutics and design. The S309 mAb, which is the parent of sotrovimab, neutralizes Omicron with 2-3-fold reduced potency compared to Wuhan-Hu-1 or Washington-1, while the 7 other clinical mAbs or mAb cocktails experience reduction of neutralizing activity of 1-2 orders of magnitude or greater. Furthermore, some Omicron isolates (≈9%) harbor the R346K substitution which in conjunction with N440K (present in the main haplotype) leads to immune escape from C135 mAb-mediated neutralization (*22, 42*). These data illustrate that mAbs targeting antigenic site IV can be distinctly affected by the Omicron mutations and underscore key differences between the strategies used to isolate S309 and C135, as R346K does not affect S309 whether in isolation or in the context of the full constellation of Omicron mutations (*6, 8, 38*). Whereas C135 was identified from a SARS-CoV-2 convalescent donor (*22*), S309 was isolated from a subject who recovered from a SARS-CoV infection in 2003 (*29*), the latter strategy increased the likelihood of finding mAbs recognizing epitope that are mutationally constrained throughout sarbecovirus evolution. The identification of broadly reactive mAbs that are neutralizing multiple distinct sarbecoviruses, including SARS-CoV-2 variants, pave the way for designing vaccines eliciting broad sarbecovirus immunity (*43*–*47*). These efforts offer hope that the same strategies that contribute to solving the current pandemic will prepare us for future putative sarbecovirus pandemics.

## Methods

### Production of recombinant spike glycoprotein

The SARS-CoV-2 S ectodomains were produced in 200 mL cultures of Expi293F Cells (ThermoFisher Scientific) grown in suspension using Expi293 Expression Medium (ThermoFisher Scientific) at 37°C in a humidified 8% CO2 incubator rotating at 130 rpm. Cells grown to a density of 2.5 million cells per mL were transfected using the ExpiFectamine 293 Transfection Kit (ThermoFisher Scientific) and cultivated for two days at which point the supernatant was harvested. S ectodomains were purified from clarified supernatants using a Cobalt affinity column (Cytiva, HiTrap TALON crude), washing with 20 column volumes of 20 mM Tris-HCl pH 8.0 and 150 mM NaCl and eluted with a gradient of 600 mM imidazole. The S ectodomain was then concentrated using a 100 kDa centrifugal filter (Amicon Ultra 0.5 mL centrifugal filters, MilliporeSigma), residual imidazole was washed away by consecutive dilutions in the centrifugal filter unit with 20 mM Tris-HCl pH 8.0 and 150 mM NaCl, and finally concentrated to 1 mg/mL before use immediately after purification.

### CryoEM sample preparation and data collection

100 μL of 1 mg/mL SARS-CoV-2 S B.1.1.529 ectodomain was incubated with 40 μl 3.4 mg/ml S309 Fab for 10 min at 37°C in 150 mM NaCl and 20 mM Tris-HCl pH 8 and then 2.2 μL of 67 mg/mL S2L20 Fab was added and the mixture was incubated for 15 min at 37°C. Unbound Fab was then washed away with six consecutive dilutions in 400 μL of 20 mM Tris-HCl pH 8.0 and 150 mM NaCl over a 100 kDa centrifugal filter (Amicon Ultra 0.5 mL centrifugal filters, MilliporeSigma). The complex was concentrated to 3.5 mg/mL and 3 μL was immediately applied onto a freshly glow discharged 2.0/2.0 UltraFoil grid (84) (200 mesh), plunge frozen using a vitrobot MarkIV (ThermoFisher Scientific) using a blot force of −1 and 6.0 s blot time at 100% humidity and 23°C.

Data were acquired using the Leginon software (*48*) to control a FEI Titan Krios transmission electron microscope equipped with a Gatan K3 direct detector and operated at 300 kV with a Gatan Quantum GIF energy filter. The dose rate was adjusted to 3.75 counts/super-resolution pixel/s, and each movie was acquired in 75 frames of 40 ms with a pixel size of 0.843 Å and a defocus range comprised between −0.2 and −2.0 μm.

### CryoEM data processing

Movie frame alignment, estimation of the microscope contrast-transfer function parameters, particle picking and extraction (with a downsampled pixel size of 1.686 Å and box size of 256 pixels2) were carried out using Warp (*49*). Reference-free 2D classification was performed using cryoSPARC (*50*) to select well-defined particle images. 3D classification with 50 iterations each (angular sampling 7.5 □ for 25 iterations and 1.8 □ with local search for 25 iterations) were carried out using Relion without imposing symmetry (*51, 52*). 3D refinements were carried out using non-uniform refinement in cryoSPARC (*53*) before particle images were subjected to Bayesian polishing using Relion (*54*) during which particles were re-extracted with a box size of 512 Å at a pixel size of 0.843 Å. Next, 86 optics groups were defined based on the beamtilt angle used for data collection. Another round of non-uniform refinement in cryoSPARC was then performed concurrently with global and per-particle defocus refinement. For focused classification, particles were symmetry-expanded in Relion, and the particles were 3D classified in Relion without alignment using a mask that encompasses part of the NTD and the S2L20 VH/VL region, or the RBD and the S309 VH/VL region. Particles in well-formed 3D classes were then used for local refinement in cryoSPARC. Reported resolutions are based on the gold-standard Fourier shell correlation of 0.143 criterion and Fourier shell correlation curves were corrected for the effects of soft masking by high-resolution noise substitution (*55, 56*).

### CryoEM model building and analysis

UCSF Chimera (*57*) and Coot (*58*) were used to fit atomic models of S2L20, S309, and SARS-CoV-2 S (PDB 7SOB) into the cryo-EM maps. The model was then refined and rebuilt into the map using Coot (*58*), Rosetta (*59, 60*), Phenix (*61*), and ISOLDE (*62*). Model validation and analysis used MolProbity (*63*), EMRinger (*64*), Phenix (*61*), and Privateer (*65*). Figures were generated using UCSF ChimeraX (*66*).

### Monoclonal antibodies

Antibody VH and VL sequences for mAbs COV2-2130 (PDB ID 7L7E), COV2-2196 (PDB ID 7L7E, 7L7D), REGN10933 (PDB ID 6XDG), REGN10987 (PDB ID 6XDG) and ADI-58125 (PCT application WO2021207597, seq. IDs 22301 and 22311) were subcloned into heavy chain (human IgG1) and the corresponding light chain (human IgKappa, IgLambda) expression vectors respectively and produced in transiently transfected ExpiCHO-S cells (Thermo Fisher, #A29133) at 37°C and 8% CO2. Cells were transfected using ExpiFectamine. Transfected cells were supplemented 1 day after transfection with ExpiCHO Feed and ExpiFectamine CHO Enhancer. Cell culture supernatant was collected eight days after transfection and filtered through a 0.2 µm filter. Recombinant antibodies were affinity purified on an ÄKTA Xpress FPLC device using 5 mL HiTrap™ MabSelect™ PrismA columns followed by buffer exchange to Histidine buffer (20 mM Histidine, 8% sucrose, pH 6) using HiPrep 26/10 desalting columns.

Antibody VH and VL sequences for LY-CoV555, LY-CoV016, and CT-P59 were obtained from PDB IDs 7KMG, 7C01 and 7CM4, respectively and mAbs were produced as recombinant IgG1 by ATUM. S309 was produced by WuXi Biologics (China). Recombinant S304 and S309 Fabs were produced by ATUM.

### Recombinant protein production

SARS-CoV-2 RBD proteins for SPR binding assays (residues 328-531 of S protein from GenBank NC_045512.2 with N-terminal signal peptide and C-terminal thrombin cleavage site-TwinStrep-8xHis-tag) were expressed in Expi293F (Thermo Fisher Scientific) cells at 37°C and 8% CO2. Transfections were performed using the ExpiFectamine 293 Transfection Kit (Thermo Fisher Scientific). Cell culture supernatants were collected three days after transfection and supplemented with 10x PBS to a final concentration of 2.5x PBS (342.5 mM NaCl, 6.75 mM KCl and 29.75 mM phosphates). RBDs were purified using cobalt-based immobilized metal affinity chromatography followed by buffer exchange into PBS using a HiPrep 26/10 desalting column (Cytiva) for Wuhan-Hu-1 protein, or a Superdex 200 Increase 10/300 GL column (Cytiva) for Omicron protein.

SARS-CoV-2 Omicron RBD for crystallization (residues 328-531, with N-terminal signal peptide and ‘ETGT’, and C-terminal 8xHis-tag) was expressed similarly as described above in the presence of 10 µM kifunensine. Cell culture supernatant was collected three days after transfection and supplemented with 10x PBS to a final concentration of 2.5x PBS. Protein was purified using a HiTrap TALON crude cartridge followed by buffer exchange into PBS using a HiPrep 26/10 desalting column (Cytiva).

ACE2 for crystallization (residues 19-615 from Uniprot Q9BYF1 with a C-terminal thrombin cleavage site-TwinStrep-10xHis-GGG-tag, and N-terminal signal peptide) was expressed in ExpiCHO cells in the presence of 10 µM kifunensine at 37°C and 8% CO_2_. Transfection was performed using the ExpiCHO transfection kit (Thermo Fisher Scientific). Cell culture supernatant was collected eight days after transfection and supplemented to a final concentration of 80 mM Tris-HCl pH 8.0, 100 mM NaCl, and then incubated with BioLock (IBA GmbH) solution. hACE2 was purified using a 5 mL StrepTrap HP column (Cytiva) followed by size exclusion chromatography using a Superdex 200 Increase 10/300 GL column (Cytiva) pre-equilibrated in PBS.

### SPR binding measurements of IgG

SPR binding measurements were performed using a Biacore T200 instrument with a CM5 sensor chip covalently immobilized using Cytiva Human Antibody Capture Kit. Running buffer was Cytiva HBS-EP+ (pH 7.4). All measurements were performed at 25 °C. RBD analyte concentrations were 3.1, 12.5, and 50 nM, run as single-cycle kinetics. Double reference-subtracted data were fit using Biacore T200 Evaluation software (version 3.1) to a 1:1 binding model except two datasets (LY-CoV016 and REGN10987 binding to Wuhan-Hu-1 RBD) were fit to a Heterogeneous Ligand model (“biphasic fit”) due to an artefactual kinetic phase with very slow dissociation that often arises when RBD is an analyte. For the Heterogeneous Ligand fits, the lower affinity of the two K_D_ values reported by the fit is taken as the K_D_ (the two K_D_ values are separated by at least two orders of magnitude). Omicron binding data are all n=1, except S309:Omicron data is n=3. Wuhan-Hu-1 RBD binding data are representative of at least n=2 for S309, CT-P59, both LY mAbs, and both REGN mAbs; others are n=1.

### Crystallization, data collection, structure determination, and analysis

Prior to forming the SARS-CoV-2 Omicron RBD-ACE2-S304-S309 complex, recombinant SARS-CoV-2 Omicron RBD was digested with EndoH over night (New England Biolabs). Recombinant hACE2 protein was digested using EndoH (New England Biolabs) and thrombin (Sigma-Aldrich) over night. RBD was mixed with a 1.3-fold molar excess of deglycosylated hACE2, S304 Fab, and S309 Fab. The complex was purified on a Superdex 200 10/300 GL column pre-equilibrated with 20 mM Tris-HCl pH 7.5, 150 mM NaCl. Crystals of the SARS-CoV-2 Omicron RBD-hACE2-S304-S309 complex were obtained at 20°C by sitting drop vapor diffusion. A total of 200 nL of the complex at 6 mg/mL were mixed with 200 nL mother liquor solution containing 0.1 M carboxylic acids (Molecular Dimensions), 20% v/v glycerol, 10% w/v PEG 4000, and 0.1 M Tris (base)/bicine pH 8.5.

Data were collected at the Molecular Biology Consortium beamline 4.2.2 at the Advanced Light Source synchrotron facility in Berkeley, CA and processed with the XDS software package for a final dataset of 3.01 Å in space group P2_1_. The SARS-CoV-2 Omicron RBD-hACE2-S304-S309 complex structure was solved by molecular replacement using Phaser from starting models consisting of RBD-S304-S309 (PDB: <u>7JX3</u>) and hACE2 (PDB: <u>6m0j</u>). Several subsequent rounds of model building and refinement were performed using Coot, ISOLDE, Refmac5, Phenix and MOE (https://www.chemcomp.com), to arrive at a final model for the quaternary complex.

## Acknowledgements

This study was supported by the National Institute of Allergy and Infectious Diseases (DP1AI158186 and HHSN272201700059C to D.V.), a Pew Biomedical Scholars Award (D.V.), an Investigators in the Pathogenesis of Infectious Disease Awards from the Burroughs Wellcome Fund (D.V.), Fast Grants (D.V.), the National Institute of General Medical Sciences (5T32GM008268-32 to SKZ), the University of Washington Arnold and Mabel Beckman cryoEM center and the National Institute of Health grant S10OD032290 (to D.V.). D.V. is an Investigator of the Howard Hughes Medical Institute. Beamline 4.2.2 of the Advanced Light Source, a U.S. DOE Office of Science User Facility under Contract No. DE-AC02-05CH11231, is supported in part by the ALS-ENABLE program funded by the National Institutes of Health, National Institute of General Medical Sciences, grant P30 GM124169-01. This research was funded in whole, or in part, by the Wellcome Trust [209407/Z/17/Z]. For the purpose of open access, the author has applied a CC BY public copyright licence to any Author Accepted Manuscript version arising from this submission.

## Author contributions

M.M., J.E.B., H.W.V., D.C., G.S. and D.V. conceived the project. M.M., L.E.R., S.K.Z., G.S. and D.V designed experiments. M.M., N.C., S.K.Z, J.R.D and A.E.P expressed and purified proteins. L.E.R. performed SPR analysis. S.K.Z. performed BLI analysis. M.M. carried out cryoEM sample preparation, data collection, processing model building and refinement. NC carried out crystallization experiments. J.N. collected and processed X-ray diffraction data. M.M., T.C., G.S. and D.V. built and refined the crystal structure. M.M. and D.V. wrote an initial draft of the manuscript with input from all authors.

## Data and Materials Availability

The cryoEM map and coordinates have been deposited to the Electron Microscopy Databank and Protein Data Bank with accession numbers TBD. The crystal structure has been deposited to the Protein Data Bank with accession number TBD.

---

**Figure S1.**
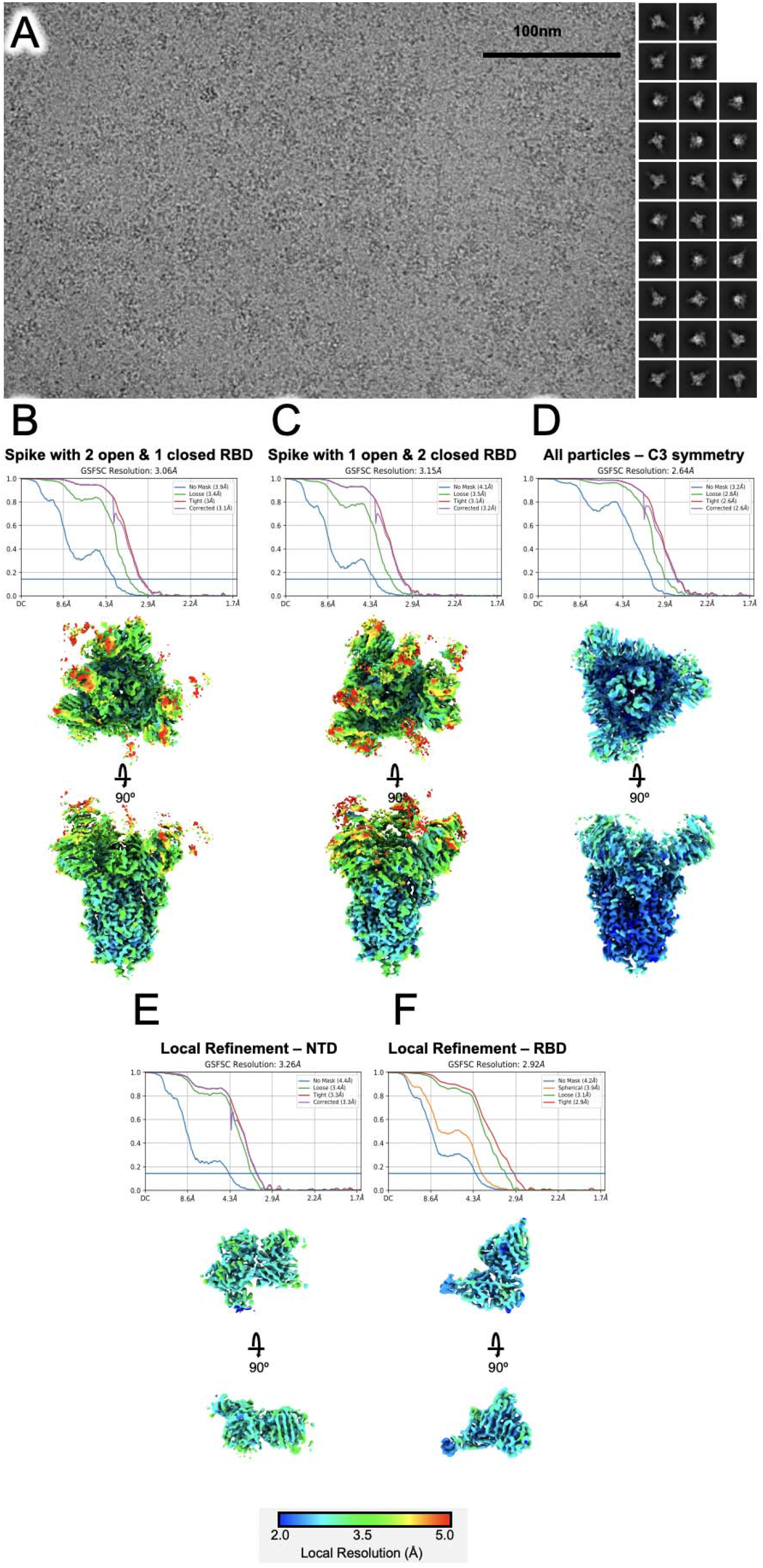
CryoEM data processing and validation. **(A)** Representative electron micrograph (left, scale bar: 100 nm) and 2D class averages (right) are shown for the indicated particles embedded in vitreous ice. (**B, C, D, E, and F**) Gold-standard Fourier shell correlation curves with the 0.143 cutoff indicated by a horizontal blue line (top) and unsharpened maps colored by local resolution calculated using cryoSPARC (bottom) for whole reconstructions (B, C, and D) and the locally refined reconstructions of NTD- or RBD-bound Fab variable domains (E and F).

**Figure S2.**
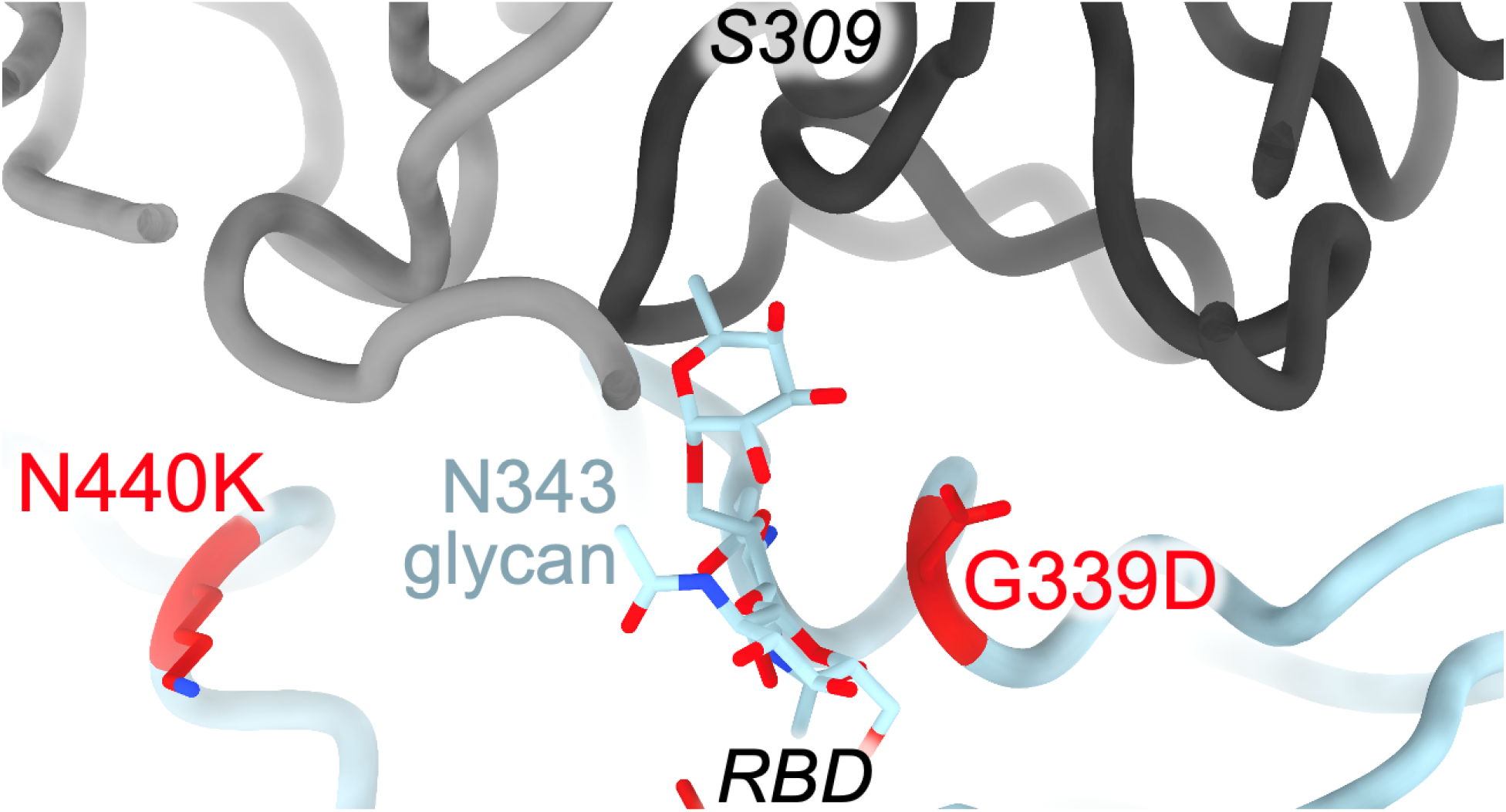
Structural basis for S309 binding to the Omicron RBD. Zoomed-in view of the cryoEM structure of the S309 Fab fragment (black/grey) bound to the Omicron RBD (blue ribbon) with mutated residues shown in red as sticks. The N343 glycan is shown as sticks.

**Fig S3.**
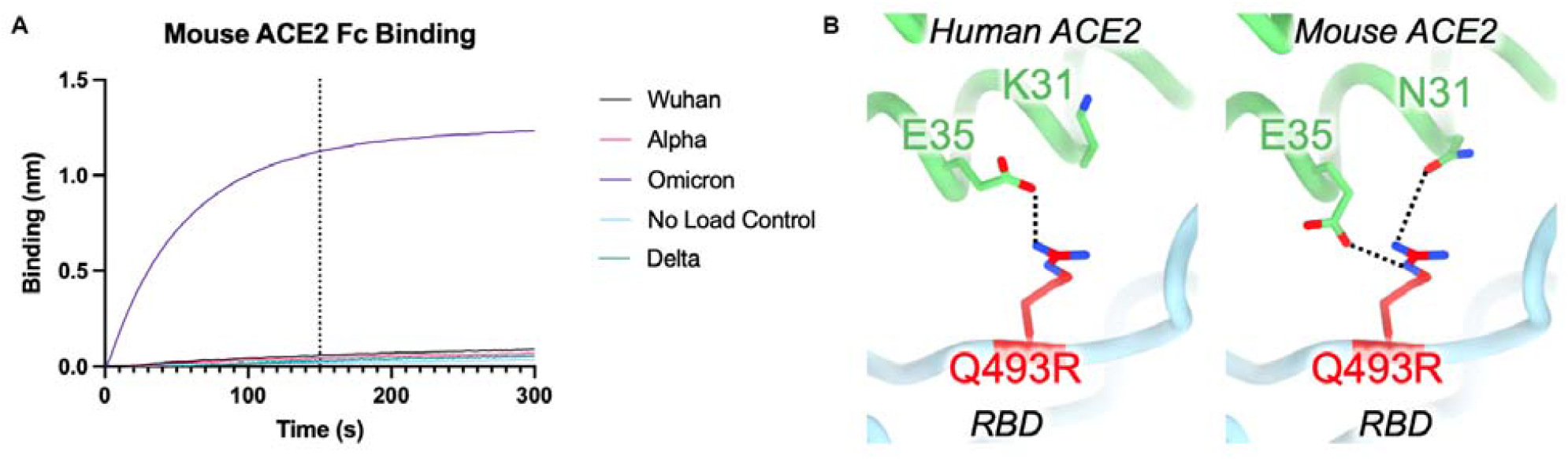
The SARS-CoV-2 Omicron RBD recognizes mouse ACE2 efficiently. **A**, Biolayer interferometry binding analysis of 1µM dimeric mouse ACE2-Fc to biotinylated SARS-CoV-2 variant RBDs immobilized at the surface of SA biosensors. The vertical dashed lines indicate the transition between association and dissociation phases. **B**, In silico model of the interface between the Omicron RBD and mouse ACE2 (right) based on the crystal structure of the human ACE2-bound Omicron RBD (left).

**Table S1:** Kinetic parameters and fitting details of the SPR binding measurements.

| IgG | RBD | ka (1/Ms) | kd (1/s) | KD (M) | Rmax (RU) | Capture level (RU) |
| --- | --- | --- | --- | --- | --- | --- |
| S309 | Wu-Hu-1 | 1.49E+05 | 1.32E-04 | 8.87E-10 | 73.77 | 210.5 |
| S309 | Omicron | 7.15E+04 | 1.90E-04 | 2.67E-09 | 82.01 | 211.8 |
| ADI-58125 | Wu-Hu-1 | 1.15E+06 | 7.17E-08<sup>a</sup> | < 1E-11<sup>a</sup> | 113.6 | 231 |
| ADI-58125 | Omicron | 5.42E+06 | 0.2497 | 4.61E-08 | 83.81 | 230.3 |
| COV2-2130 | Wu-Hu-1 | 4.26E+05 | 8.38E-04 | 1.97E-09 | 109.5 | 251.7 |
| COV2-2130 | Omicron | 9.30E+05 | 0.05897 | 6.34E-08 | 79.31 | 246.7 |
| COV2-2196 | Wu-Hu-1 | 1.14E+07 | 0.01244 | 1.09E-09 | 100.7 | 240.5 |
| COV2-2196 | Omicron | 9.04E+05 | 0.441 | 4.88E-07 | 100<sup>b</sup> | 239.7 |
| CT-P59 | Wu-Hu-1 | 5.54E+06 | 3.65E-04 | 6.60E-11 | 97.99 | 236.8 |
| CT-P59 | Omicron | No binding |  |  |  | 236.9 |
| LY-CoV016 | Wu-Hu-1 | See Biphasic fit results, below |  |  |  | 196.1 |
| LY-CoV016 | Omicron | No binding |  |  |  | 195 |
| LY-CoV555 | Wu-Hu-1 | 7.92E+05 | 0.001174 | 1.48E-09 | 100.7 | 242.1 |
| LY-CoV555 | Omicron | No binding |  |  |  | 243.6 |
| REGN10933 | Wu-Hu-1 | 5.29E+06 | 0.003645 | 6.89E-10 | 93.53 | 260 |
| REGN10933 | Omicron | 6.16E+05 | 0.5073 | 8.24E-07 | 94<sup>b</sup> | 259.6 |
| REGN10987 | Wu-Hu-1 | See Biphasic fit results, below |  |  |  | 231.6 |
| REGN10987 | Omicron | No binding |  |  |  | 231.5 |

<sup>a</sup>Measured dissociation is beyond the limit of detection. Reporting KD as an upper bound
<sup>b</sup>Rmax fixed to Rmax from WT fit (without constraint, Rmax from Omicron fit was implausibly much lower than Rmax for corresponding Wuhan-Hu-1 binding)

Biphasic fit (Heterogeneous Ligand Model), binding to Wuhan-Hu-1 RBD Kinetic phase attributed to real binding (vs an artifact) highlighted in bold
| IgG | ka1<br>(1/Ms) | kd1<br>(1/s) | KD1 (M) | ka2<br>(1/Ms) | kd2 (1/s) | KD2 (M) | Rmax1<br>(RU) | Rmax2<br>(RU) |
| --- | --- | --- | --- | --- | --- | --- | --- | --- |
| LY-CoV016 | **1.24E+06** | **0.0172** | **1.38E-08** | 1.79E+05 | 5.23E-05 | 2.91E-10 | **59.66** | 20.68 |
| REGN10987 | **1.28E+06** | **0.0159** | **1.24E-08** | 2.74E+05 | 1.02E-08 | 3.73E-14 | **70.48** | 28.6 |

**Table S2:** Crystallographic data collection and refinement statistics.

|  | Omicron RBD/<br>hACE2/S309/S304 |
| --- | --- |
| **Data collection** |  |
| Space group | P2<sub>1</sub> |
| Cell dimensions |  |
| *a*, *b*, *c* (Å) | 78.82, 185.82, 193.96 |
| $\alpha$, $\beta$, $\gamma$ (°) | 90.00, 96.05, 90.00 |
| Resolution (Å) | 48.60-3.01 (3.06-3.01) |
| *R*<sub>merge</sub> | 0.539 (1.807) |
| *I* / $\sigma I$ | 3.2 (0.8) |
| Completeness (%) | 98.5 (99.3) |
| Redundancy | 3.7 (3.5) |
| **Refinement** |  |
| Resolution (Å) | 3.01 |
| No. reflections | 106,557 |
| *R*<sub>work</sub> / *R*<sub>free</sub> | 0.31/0.33 |
| No. atoms |  |
| Protein | 25,481 |
| Ligand/ion | 135 |
| Water | 63 |
| *B*-factors |  |
| Protein | 39.91 |
| Ligand/ion | 41.00 |
| Water | 41.76 |
| R.m.s. deviations |  |
| Bond lengths (Å) | 0.004 |
| Bond angles (°) | 0.815 |

\*Values in parentheses are for highest-resolution shell.

